# Localizing post-admixture adaptive variants with object detection on ancestry-painted chromosomes

**DOI:** 10.1101/2022.09.04.506532

**Authors:** Iman Hamid, Katharine L. Korunes, Daniel R. Schrider, Amy Goldberg

**Affiliations:** Department of Evolutionary Anthropology, Duke University, Durham, North Carolina, USA; Department of Genetics, University of North Carolina, Chapel Hill, North Carolina, USA

## Abstract

Gene flow between previously isolated populations during the founding of an admixed or hybrid population has the potential to introduce adaptive alleles into the new population. If the adaptive allele is common in one source population, but not the other, then as the adaptive allele rises in frequency in the admixed population, genetic ancestry from the source containing the adaptive allele will increase nearby as well. Patterns of genetic ancestry have therefore been used to identify post-admixture positive selection in humans and other animals, including examples in immunity, metabolism, and animal coloration. A common method identifies regions of the genome that have local ancestry ‘outliers’ compared to the distribution across the rest of the genome, considering each locus independently. However, we lack theoretical models for expected distributions of ancestry under various demographic scenarios, resulting in potential false positives and false negatives. Further, ancestry patterns between distant sites are often not independent. As a result, current methods tend to infer wide genomic regions containing many genes as under selection, limiting biological interpretation. Instead, we develop a deep learning object detection method applied to images generated from local ancestry-painted genomes. This approach preserves information from the surrounding genomic context and avoids potential pitfalls of user-defined summary statistics. We find the-method is robust to a variety of demographic misspecifications using simulated data. Applied to human genotype data from Cabo Verde, we localize a known adaptive locus to a single narrow region compared to multiple or long windows obtained using two other ancestry-based methods.

## Introduction

Genetic exchange between previously separated populations is ubiquitous across species (Moran et al., 2021; Payseur & Rieseberg 2016), often referred to as ‘admixture’ or ‘hybridization’ when moderate - to large-scale movements of individuals create new populations with ancestors from multiple source populations. In admixed populations, genetic ancestry varies between individuals and along the chromosome within individuals (Aguillon et al., 2022; Gopalan et al., 2022; Hellenthal et al., 2014). Across the tree of life, variation in genetic ancestry shapes genetic and phenotypic variation, such as differences in disease risk between populations. Small amounts of gene flow or larger admixture may introduce advantageous alleles which then undergo positive selection. Such cases have been identified in diverse taxa, often termed adaptive introgression (Aguillon et al., 2022; Edelman & Mallet 2021; Hedrick, 2013; Hsieh et al., 2019; Huerta-Sánchez et al., 2014; Moran et al., 2021; Norris et al., 2015; Oziolor et al., 2019; Racimo et al., 2015; Whitney et al., 2006) or, in humans, post-admixture positive selection (Cuadros-Espinoza et al., 2022; Gopalan et al., 2022; Tang et al., 2007).

Despite the ubiquity and biological importance of admixture, understanding evolutionary processes in admixed populations remains challenging (Gopalan et al., 2022; Moran et al., 2021). Classical methods to detect selection may pick up signatures of pre-admixture selection, and are often confounded by the process of admixture, which can increase linkage disequilibrium (LD) and change the distribution of allele frequencies (Cuadros-Espinoza et al., 2022; Lohmueller et al., 2010, 2011; Yelman et al., 2021). Yet, because admixture can introduce advantageous alleles at intermediate frequencies, post-admixture selection provides an opportunity for particularly rapid adaptation on the scale of tens or hundreds of generations (Hellenthal et al., 2016; Hamid et al., 2021). Thus, methods tailored to the genetic signatures of admixed populations are important to investigate the extent and impact of post-admixture adaptation across many organisms.

Recent methods have advanced our ability to identify regions of admixed genomes containing haplotypes under positive selection by using patterns of genetic ancestry. When one source population provides a beneficial allele, we expect that, as the beneficial allele increases in frequency, linked alleles from the source population will hitchhike along with it, and thereby the proportion of admixed individuals with ancestry from that source population at the selected locus (i.e. the local ancestry proportion) increases too. This logic has been leveraged to detect selection in recently admixed populations by identifying outliers in local ancestry proportion compared to a genome-wide average. Applied to human populations, variations on ancestry outlier detection have identified genomic regions associated with a range of phenotypic traits potentially underlying adaptation, including response to high altitude, diet, pigmentation, immunity, and disease susceptibility (Bryc et al., 2010; Bryc et al., 2015; Busby et al., 2016; Busby et al., 2017; Cuadros-Espinoza et al., 2022; Fernandes et al., 2019; Hamid et al., 2021; Isshiki et al., 2021; Jeong et al., 2014; Jin et al., 2012; Laso-Jadart et al., 2017; Lopez et al., 2019; Norris et al., 2020; Patin et al., 2017; Pierron et al., 2018; Rishishwar et al., 2015; Tang et al., 2007; Triska et al., 2015; Vicuña et al., 2020; Zhou et al., 2016).

This ancestry outlier detection approach is useful for identifying regions that may be under selection, but it can yield false positives due to long-range LD from the source populations or allele frequencies drifting as a result of serial founder effects, and the criteria for determining outliers is difficult (Bhatia et al., 2014; Buby et al., 2017; Price et al., 2008); false negatives may also occur if the number of true adaptive events is greater than the number of outliers retained. Importantly, the ancestry outlier approach discards the wealth of information from the surrounding genomic context. Along-genome spatial patterns of ancestry, such as the distribution of ancestry tract lengths containing a selected locus, may be informative about selection on this timescale in admixed populations. The length of ancestry tracts is influenced by the timing and strength of selection, analogous to the increase in LD around selective sweeps in homogeneous populations (Kelley 1997; Kim & Nielsen, 2004; Sabeti et al., 2002; Voight et al., 2006). Similarly, strong selection can influence ancestry patterns along long stretches of the genome, often in complex patterns depending on the evolutionary scenario (Hamid et al., 2021; Shchur et al., 2020; Svedberg et al., 2021). For example, Svedberg et al. 2021 extend their prior model (Ancestry_HMM, Corbett-Detig & Nielsen 2017) to explicitly incorporate post-admixture selection by modeling increased ancestry frequency at the selected allele and a longer introgressed haplotype. We used similar expected signatures summarized in the *iDAT* statistic developed in Hamid et al. 2021. However, the expected distributions of the length and frequency of ancestry tracts surrounding post-admixture positively selected alleles has been difficult to explore theoretically, particularly combined with variable demographic histories (with the notable exception of Shchur et al. 2020).

However, information about the complex patterns of ancestry around a selected locus is lost when relying on summary statistics, and there is a bias inherent in the user’s choice of quantitative summaries to include during inference. More generally, we lack theoretical expectations for patterns of ancestry expected under post-admixture selection, especially under a range of selective and demographic histories.

To overcome the loss of spatial information along the genome and the simplifying assumptions of classical summary statistics, deep learning techniques have been increasingly used in population genetics. Deep learning algorithms are multi-layered networks trained on example datasets with known response variables with the goal of learning a relationship between the input data and output variable(s) (applications to population genetics reviewed in Schrider & Kern (2018). Deep learning techniques are flexible with respect to data type and the specific task at hand, and have been shown to be effective for inferring demographic histories (Flagel et al., 2019; Sanchez et al., 2021; Sheehan & Song, 2016; Wang et al., 2021), recombination rates (Adrion et al., 2020; Chan et al., 2018; Flagel et al., 2019), and natural selection (Gower et al., 2021; Kern & Schrider, 2018; Sheehan & Song, 2016). Among the branches of deep learning, computer vision methods are a family of techniques originally developed to recognize images by using convolutional neural networks (CNNs) (Krizhevsky et al., 2012; LeCun et al., 2015; Lecun & Bengio, 1995). CNNs learn from complex spatial patterns in large datasets through a series of filtering and down sampling operations that compress the data into features that are informative for inference. CNNs have recently been applied to images of genotype matrices for population genetic inference with great success (Battey et al., 2020; Battey et al., 2021; Blischak et al., 2021; Chan et al., 2018; Flagel et al., 2019; Gower et al., 2021; Isildak et al., 2021; Sanchez et al., 2021; Torada et al., 2019). In doing so, researchers can circumvent the loss of information and bias from using user-defined population genetic summary statistics and make inferences for study systems and questions for which we lack theoretical expectations. Simulation-based inference is also often flexible enough that one may be able to incorporate various demographic histories into models, which has proven difficult for theoretical models.

Here, we build on recent successes in deep learning applications to population genetics problems and develop a deep learning object detection strategy that localizes genomic regions under selection from images of chromosomes ‘painted’ by ancestry (Figure 1) (Lawson et al., 2012; Maples et al., 2012). In using local ancestry rather than the genotypes directly, we focus on post-admixture processes and are potentially well-suited to low coverage or sparse SNP data common in non-model systems (Schaefer et al., 2016; Schaefer et al., 2017; Schumer et al., 2020; Wall et al., 2016). Using this approach, we demonstrate that complex ancestry patterns beyond single-locus summary statistics are informative about selection in recently admixed populations. We take advantage of existing deep learning object detection frameworks, illustrating the ease of use and accessibility of deep learning applications for population genetic researchers without experience in machine learning techniques. In simulated as well as human SNP data, we show that our method is able to localize regions under positive selection post admixture, and remains effective at identifying selection under a range of demographic misspecifications. We focus on scenarios with moderate to high admixture contributions occurring in the last tens to hundreds of generations; multiple other methods have recently been developed focused on older admixture scenarios at low admixture contribution rates, often termed adaptive introgression (Gower et al., 2021; Racimo et al., 2017; Setter et al., 2020; Svedberg et al., 2021).

**Figure 1.**
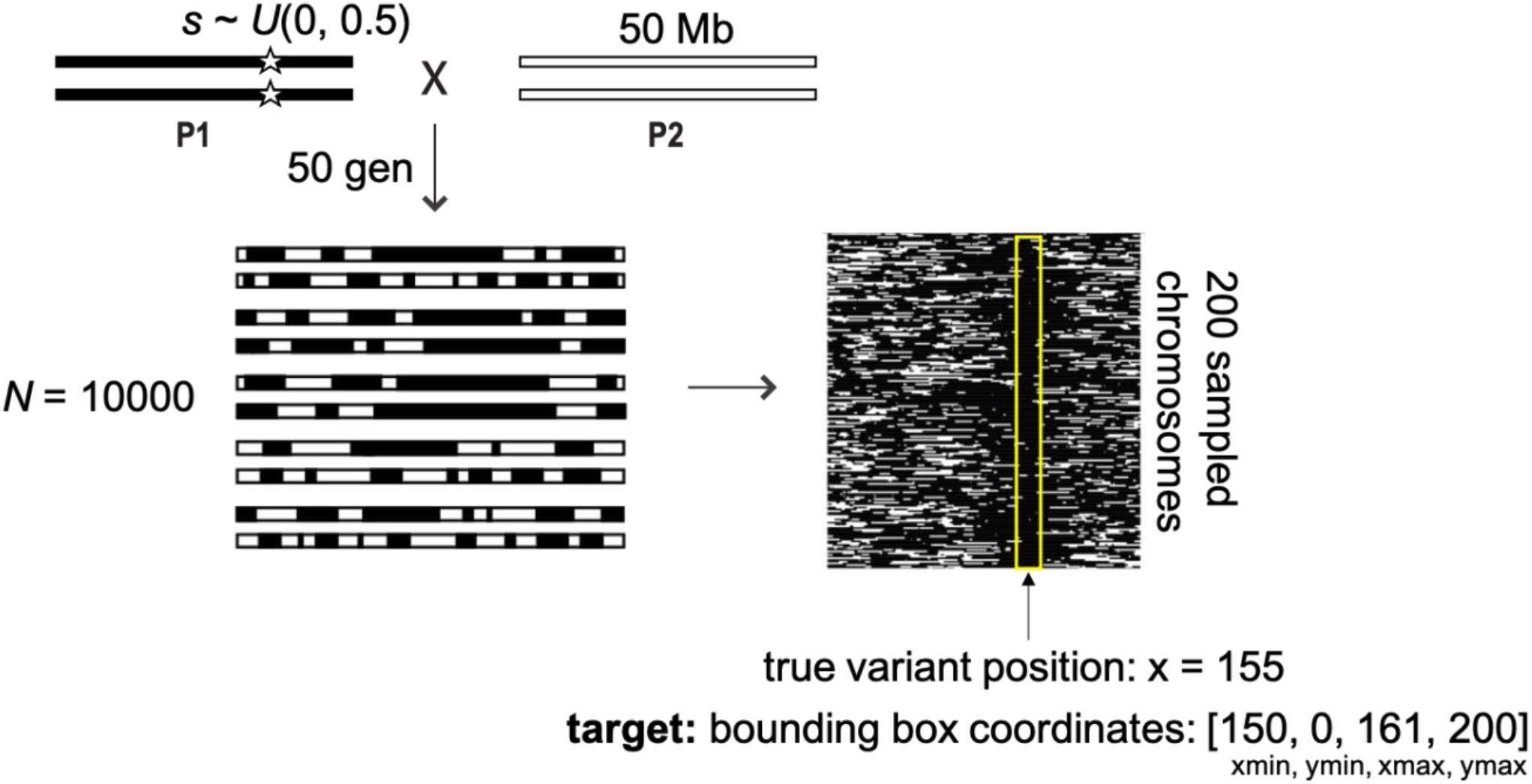
Schematic of our baseline simulation scenario. Image input for the object detection model is generated by sampling 200 ancestry-painted chromosomes from a simulated admixed population. Rows represent individuals, with chromosome position along columns. Training samples have a known “target” bounding box (yellow box), spanning an 11-pixel window centered on the position of the known beneficial variant. Using training examples, the object detection model learns the complex patterns of ancestry indicative of positive selection post-admixture and uses this information to localize a beneficial variant to a small genomic region. The trained object detection model is then expected to output bounding boxes that contain variants under selection.

## Results

### Baseline Model Performance

We first describe the object detection method’s performance in a baseline simulated scenario, before exploring the effects of model misspecification and finally comparing the method to other approaches. Full details on simulations, image generation, model training, and performance metrics are in Materials and Methods. Briefly, in the baseline scenario, we simulated a single-pulse admixture event between two isolated source populations. One source population was fixed for a beneficial variant randomly placed along the 50 Mb chromosome tract, with positive selection strength post admixture drawn from a uniform distribution *s ~ U*(0, 0.5). For each simulation, we generated two images representing two types of genetic data that a user may be analyzing: one with full local ancestry (the high resolution scenario) representing whole-genome, high-density SNP, or similar data, and the second scenario with only 100 ancestry informative markers (AIMs, the low resolution scenario) in the 50 Mb. We then trained and validated the method for each of these two sets of images. Performance metrics included precision and recall (P-R), the proportion of inferred bounding boxes that contain the true selected variant, the average width of the inferred bounding boxes, and the average number of inferred bounding boxes per image.

Overall, the locus simulated to be under positive selection was contained within the inferred bounding box ~95% of the time in both the full ancestry (high resolution) and low-resolution scenarios (Table 1 & Figure 2). As expected, the high-resolution ancestry scenario had higher precision and recall across the range of detection thresholds (Figure 2), though both had P-R curves well above a no-skill (random) classifier.

**Table 1.** Performance of object detection method on images with high and low ancestry resolution.

| ancestry resolution | bbox detection rate | average width | average number of bounding boxes | precision | recall | AUC |
| --- | --- | --- | --- | --- | --- | --- |
| high (full ancestry) | 0.950 | 10.830<br>(var = 0.615,<br>n = 1978) | 1.027<br>(var = 0.063,<br>n = 2000) | 0.886 | 0.897 | 0.871 |
| low (100 AIMs) | 0.950 | 10.834<br>(var = 0.580,<br>n = 1964) | 1.0175<br>(var = 0.064,<br>n = 2000) | 0.867 | 0.870 | 0.811 |

**Figure 2.**
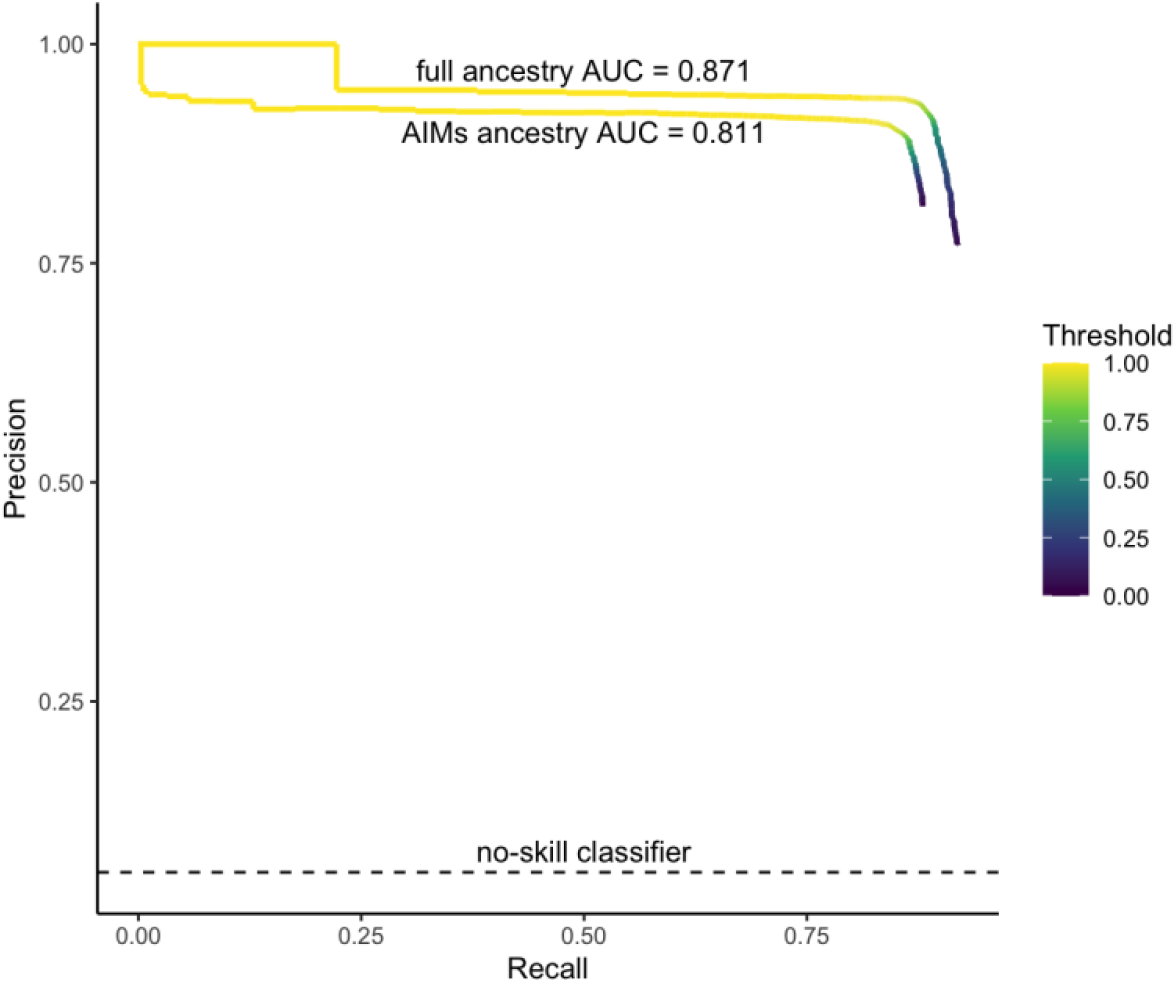
Precision-Recall curves for high (full) and low (AIMs) ancestry resolution images across a range of detection thresholds. Area under the curve (AUC) is calculated for the two scenarios, with the no-skill classifier indicated by the dashed black line.

### Model Misspecification

Often, we do not know the full model and parameters of a population’s history. We tested the robustness of our method to several demographic model misspecifications, performing inference based on images generated from simulations that differed in model and/or parameter from the ones used to generate training images. Generally, we followed the high-resolution full ancestry baseline scenario described above and in the Materials and Methods, and altered one aspect of the admixed population’s history for each scenario. We separately altered parameters for the admixture proportion, the number of generations since admixture occurred, as well as different models of the population size trajectory (bottleneck with a return to original size, expansion, or contraction). We also considered a scenario in which both source populations have the beneficial mutation segregating at a frequency of 0.5 at the time of admixture (i.e. *F_ST_* = 0 between the source populations at this allele) (see also Gopalan et al., 2022 for post-admixture positive selection simulations under different *F_ST_* values between sources at the adaptive locus)

That is, we trained the model once under the baseline scenario, and then conducted inference on simulated versions that represent empirical data under different evolutionary scenarios. We then evaluated performance using the same set of metrics as described above for the baseline model, presented in Table 2 & Figure S1. Under these demographic misspecifications, the model was still able to detect 80-98% of variants under selection, except in two scenarios where the impact of selection on patterns of local ancestry is expected to be very weak or entirely absent (Table 2).

**Table 2.**
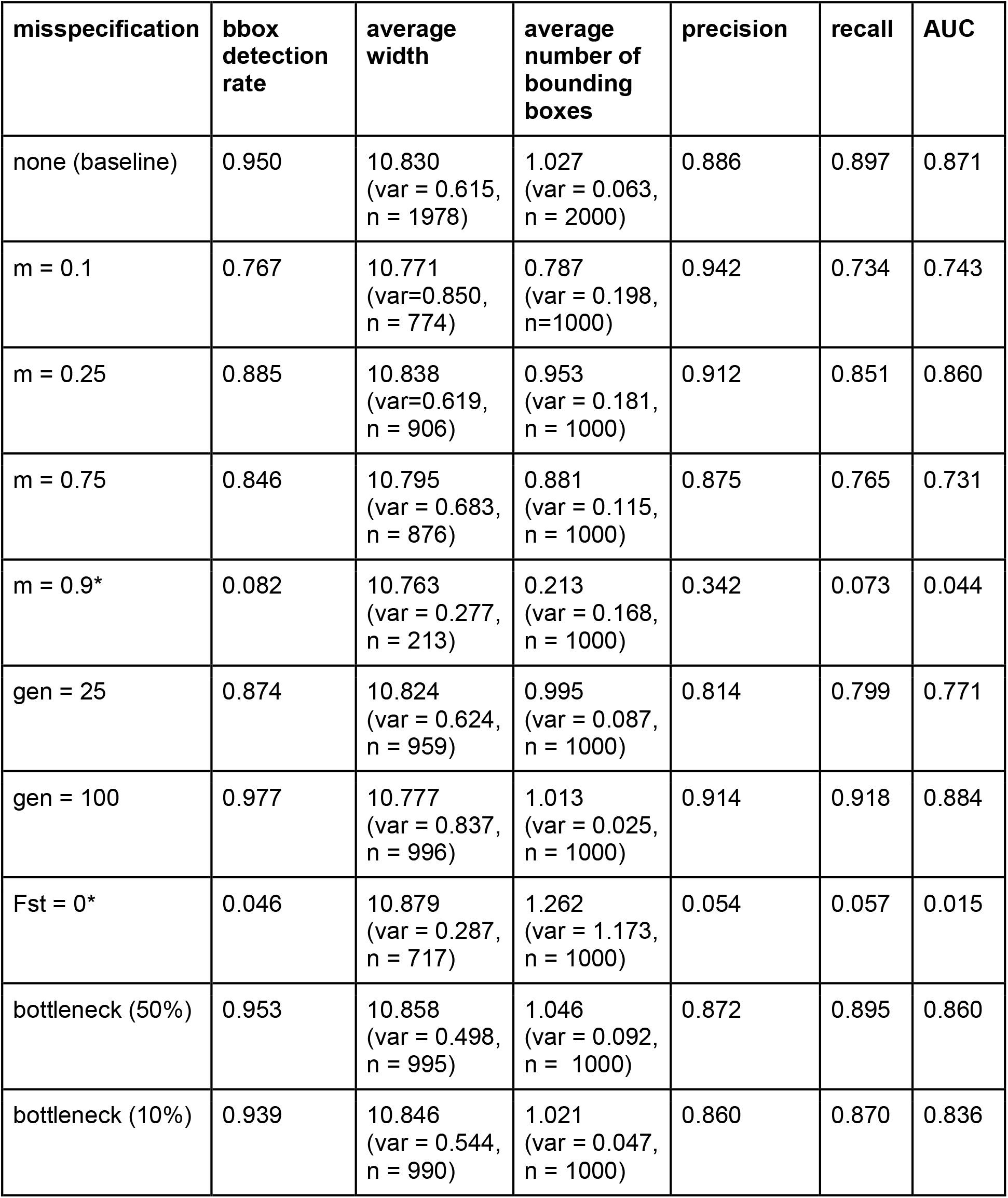
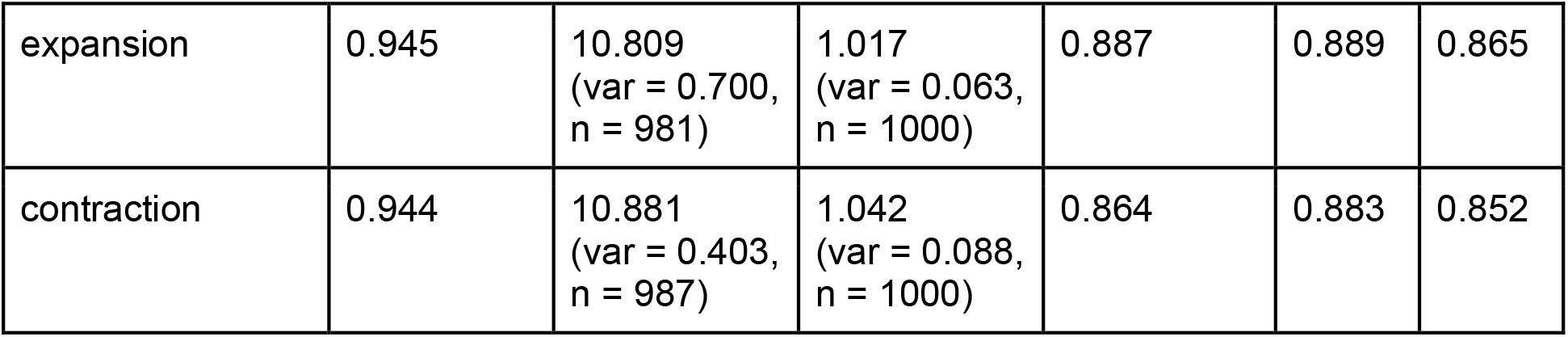
Performance of object detection method on images generated from demographic misspecifications. Further details of models in Materials and Methods, Figure S1. The two scenarios that perform poorly are marked (*).

| misspecification | bbbox<br>detection<br>rate | average<br>width | average<br>number of<br>bounding<br>boxes | precision | recall | AUC |
| --- | --- | --- | --- | --- | --- | --- |
| none (baseline) | 0.950 | 10.830<br>(var = 0.615,<br>n = 1978) | 1.027<br>(var = 0.063,<br>n = 2000) | 0.886 | 0.897 | 0.871 |
| m = 0.1 | 0.767 | 10.771<br>(var=0.850,<br>n = 774) | 0.787<br>(var = 0.198,<br>n=1000) | 0.942 | 0.734 | 0.743 |
| m = 0.25 | 0.885 | 10.838<br>(var=0.619,<br>n = 906) | 0.953<br>(var = 0.181,<br>n = 1000) | 0.912 | 0.851 | 0.860 |
| m = 0.75 | 0.846 | 10.795<br>(var = 0.683,<br>n = 876) | 0.881<br>(var = 0.115,<br>n = 1000) | 0.875 | 0.765 | 0.731 |
| m = 0.9* | 0.082 | 10.763<br>(var = 0.277,<br>n = 213) | 0.213<br>(var = 0.168,<br>n = 1000) | 0.342 | 0.073 | 0.044 |
| gen = 25 | 0.874 | 10.824<br>(var = 0.624,<br>n = 959) | 0.995<br>(var = 0.087,<br>n = 1000) | 0.814 | 0.799 | 0.771 |
| gen = 100 | 0.977 | 10.777<br>(var = 0.837,<br>n = 996) | 1.013<br>(var = 0.025,<br>n = 1000) | 0.914 | 0.918 | 0.884 |
| Fst = 0* | 0.046 | 10.879<br>(var = 0.287,<br>n = 717) | 1.262<br>(var = 1.173,<br>n = 1000) | 0.054 | 0.057 | 0.015 |
| bottleneck (50%) | 0.953 | 10.858<br>(var = 0.498,<br>n = 995) | 1.046<br>(var = 0.092,<br>n = 1000) | 0.872 | 0.895 | 0.860 |
| bottleneck (10%) | 0.939 | 10.846<br>(var = 0.544,<br>n = 990) | 1.021<br>(var = 0.047,<br>n = 1000) | 0.860 | 0.870 | 0.836 |
| expansion | 0.945 | 10.809<br>(var = 0.700,<br>n = 981) | 1.017<br>(var = 0.063,<br>n = 1000) | 0.887 | 0.889 | 0.865 |
| contraction | 0.944 | 10.881<br>(var = 0.403,<br>n = 987) | 1.042<br>(var = 0.088,<br>n = 1000) | 0.864 | 0.883 | 0.852 |

First, the model underperforms when contributing ancestry proportions was varied such that we inferred from images generated under an admixture scenario with 90% ancestral contribution from the source population providing the beneficial allele (*m* = 0.9). In this scenario the method has difficulty detecting regions under selection resulting in a high rate of false negatives (Figure S1A) because, even in regions unaffected by selection, the image is primarily one color by the end of 50 generations. We do not see this effect in the opposite scenario involving 10% ancestral contribution from the source population providing the beneficial allele (*m* = 0.1). In this scenario, the beneficial allele increasing in frequency results in the “minor” image color increasing specifically around that region.

Second, the model also underperforms when the two source populations carry the beneficial allele at the same frequency (*F_ST_* = 0). The performance of the model under this misspecification follows the no-skill classifier (Figure S1D), suggesting the model is randomly assigning bounding boxes. In this case, the model is unable to detect any ancestry-based patterns of selection because both ancestries are being equally selected. We have previously suggested and demonstrated this same result with other ancestry-based signatures of selection (Gopalan et al., 2022; Hamid et al., 2021).

### Performance on neutrally evolving chromosomes

Thus far, we have tested performance on positive examples (i.e. simulated chromosomes with a positively selected variant); here we consider negative examples where the correct inference would be that there are no regions under selection. Our method as described above is flexible enough to infer 0, 1, or multiple bboxes. However, we did not initially provide any negative examples in our training, which may impact performance for a truly neutrally evolving chromosome. First, we test our current model performance on simulated negative examples, then we train a new model including such examples.

First, we generated 1000 full ancestry images for neutrally evolving chromosomes generated under our baseline demographic model. We performed inference using our originally trained full ancestry model without training on neutral images. At a detection threshold (“bbox score”) of 0.5, our standard setting, the model predicted no bbox for 26.5% of images (see Materials and Methods for an explanation of the detection threshold parameter). For the remaining 73.5% of images, the average bbox score is 0.660, indicating overall low confidence in the predictions. If we increase the detection threshold to a bbox score of 0.7, the model predicted no bbox for 63.2% of images. If we increase the detection threshold to a bbox score of 0.9, the model predicts no bbox for 94.9% of images. For comparison, on the original validation set, the average bbox score is 0.972. To summarize, by increasing the detection threshold, one can weed out low confidence predictions and have high accuracy on neutrally evolving chromosomes.

Next, we train our model including neutral simulations (“negative examples”) to understand the potential benefits of more tailored training sets. We trained a random subset of our original training images but included neutral images as well (training set = 800 total images [640 selection images, 160 neutral images], validation set = 200 total images [180 selection, 40 neutral]). Then, we tested the newly trained model on the remaining 800 neutral images. We find that of these, 797 (>99%) accurately predict no variant under selection (meaning no bounding boxes are predicted), while 3 (0.375%) predict a variant under selection even at a detection threshold of 0.5 (model default, but relatively low confidence). When we increased the detection threshold to 0.75 to include only high confidence predictions, 100% of the neutral simulations were correctly predicted to have no bboxes.

Accuracy on selected images (n=9180) remains high in this newly trained model with 90.8% of predicted bounding boxes containing a selected variant (precision: 0.904, recall 0.828 at a detection threshold of 0.5). This is trained on a much smaller dataset than the original model, which explains the slightly lower overall performance.

### Performance on chromosomes with multiple selected variants

We primarily considered scenarios with a single locus under selection, yet depending on the window size considered, there may be multiple sites under selection. There are many complex scenarios that one could possibly test based on combinations of the number of loci across various selection strengths at different spacing between variants. In order to gain a general intuition for the model performance in scenarios where multiple sites are hypothesized to be under selection, we consider a simple example and outline a possible solution to improve performance in similar cases.

If multiple selected sites are in close proximity, their ancestry signals may interfere with one another, and the model may have difficulty distinguishing the signals resulting in the model predicting a broad region or a region between the two sites to be under selection. If one site has undergone much stronger selection than the other, the model may only confidently identify the stronger signal. As a simple example, we generated 10 images with two sites under equal selection strengths (s=0.05 for both sites). We generated a large chromosome (250 Mb, roughly the size of human chromosome 1), and placed the selected variants near opposite ends of the chromosome so their signals would not interfere with one another; variant 1: 10% of the chromosome length (physical position = 25 Mb); variant 2: 90% of the chromosome length (225 Mb). Both variants were fixed in ancestral population 1 and absent in ancestral population 2, so that the selection signal would come from the same ancestry for both sites. The demographic scenario followed our baseline trained model. The model, which was trained with a single positively-selected locus, correctly picked out at least one selected variant for 10 out of 10 images. The model was able to identify both selected variants for 5 out of the 10 images.

Alternatively, if one wanted to use the model pre-trained with a single selected locus, and reasonably suspected multiple sites were under selection, one could consider splitting large chromosomes into smaller chunks in order to pick up multiple sites. To test this scenario, we split the 10 chromosomes from the example above in half to generate two separate images, each containing only one selected variant. In this case, the model was able to detect the selected variants for 100% of images.

### Comparison to ancestry outlier detection

We next sought to evaluate whether our method constitutes an improvement on the most commonly used method for detecting regions under selection for admixed populations.The ‘local ancestry outlier’ approach identifies regions that deviate from the genome-wide average ancestry proportion, which are hypothesized to be enriched for regions under selection (Bryc et al., 2010; Gopalan et al., 2022; Tang et al., 2007). We compared performance between ancestry outlier detection and our method by calculating precision and recall, including over a range of selection coefficients (Table 3 & Figure 3B-E). For each genomic window, we additionally calculated the proportion of simulations that were classified as being “under selection” at that region as a measure of localization ability (Figure 3A). The local ancestry approach has much lower precision resulting from increased false positives, even in scenarios with greater selection strength (Table 3 & Figure 3B&C). This is further visualized in Figure 3A, where the object detection method detects a narrower region under selection (~3 Mb) compared to the local ancestry outlier approach (~8 Mb). The width of the inferred region in object detection is highly determined by the bbox size in training data, as well as window length and input image size so it is likely possible to narrow the inferred region further.

**Table 3.** Performance of object detection and local ancestry outlier methods.

| method | bbox detection rate | average width | average number of bounding boxes | precision | recall |
| --- | --- | --- | --- | --- | --- |
| object detection | 0.948 | 10.990<br>(var = 0.002,<br>n = 990) | 1.037<br>(var = 0.092,<br>n = 1000) | 0.875 | 0.890 |
| local ancestry outlier | - | - | - | 0.542 | 0.901 |

**Figure 3.**
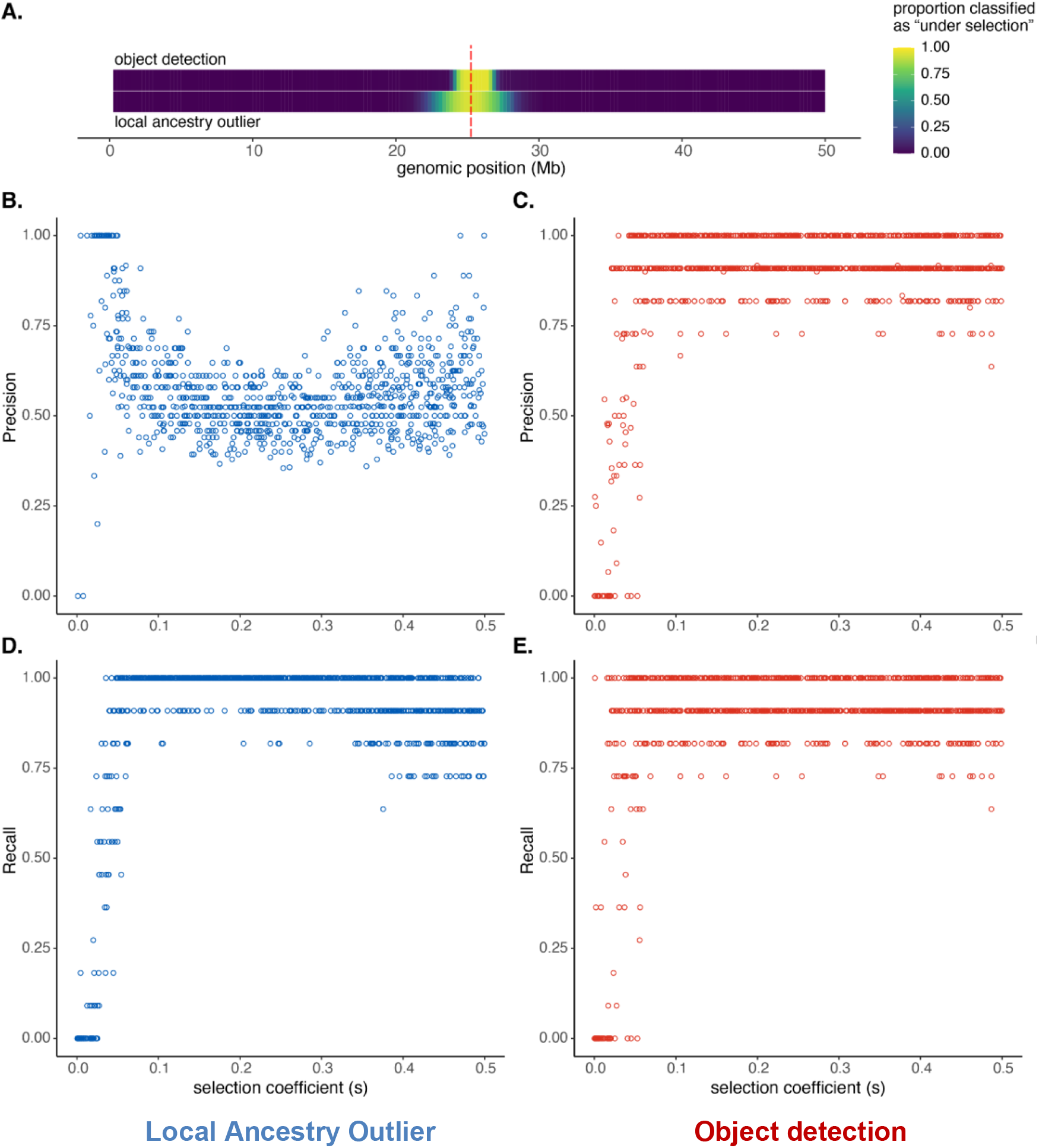
Comparison of local ancestry outlier approach and object detection method. A) Heatmap showing, for each genomic window, the proportion of simulations that had that region classified as “under selection” by either the object detection (top) or local ancestry outlier (bottom) methods. The position of the true selected variant is indicated by the vertical dashed red line. Precision across a range of selection coefficients (*s*) for the B) local ancestry outlier approach and C) the object detection method. Recall across a range of selection coefficients (*s*) for the D) local ancestry outlier approach and E) object detection method. (Also see Figures S2 and S3.)

### Application to human genotype data from Cabo Verde

We next tested the object detection method on human genotype data from the admixed population of Santiago, Cabo Verde using genotype data from 172 individuals at ~800k SNPs genome-wide (Beleza et al., 2013). We previously showed multiple lines of evidence for adaptation in this dataset at the Duffy-null that is protective against *P. vivax* malaria, including ancestry outlier detection and a statistic that incorporates the length of tracts as well as their frequency, *iDAT;* this allele is common in African ancestry and rare in Portuguese ancestry (Hamid et al., 2021). This locus has been a candidate for post-admixture positive selection in multiple other populations as well (Busby et al., 2017; Fernandes et al., 2019; Hodgson et al., 2014; Laso-Jadart et al., 2017; Pierron et al., 2018; Triska et al., 2015).

We test for post-admixture selection along the entirety of chromosome 1. Figure 4 shows that all three methods detect an adaptive locus in the nearby region; the object detection approach is highly specific, returning a single bbox approximately centered on the adaptive locus (center is ~130 kb from truth), whereas the ancestry-outlier approach returns multiple nearby hits across ~48 Mb (outliers sum to ~6 Mb). iDAT finds one region as an outlier spanning ~12 Mb and not centered on the locus under selection. The nearby centromere may be extending the window that ancestry outlier detection identifies as under selection by repressing recombination. We generated the image of ancestry on Santiago using genetic distances so the object detection approach is less sensitive to recombination variation without needing to explicitly model recombination variation in the training data.

**Figure 4.**
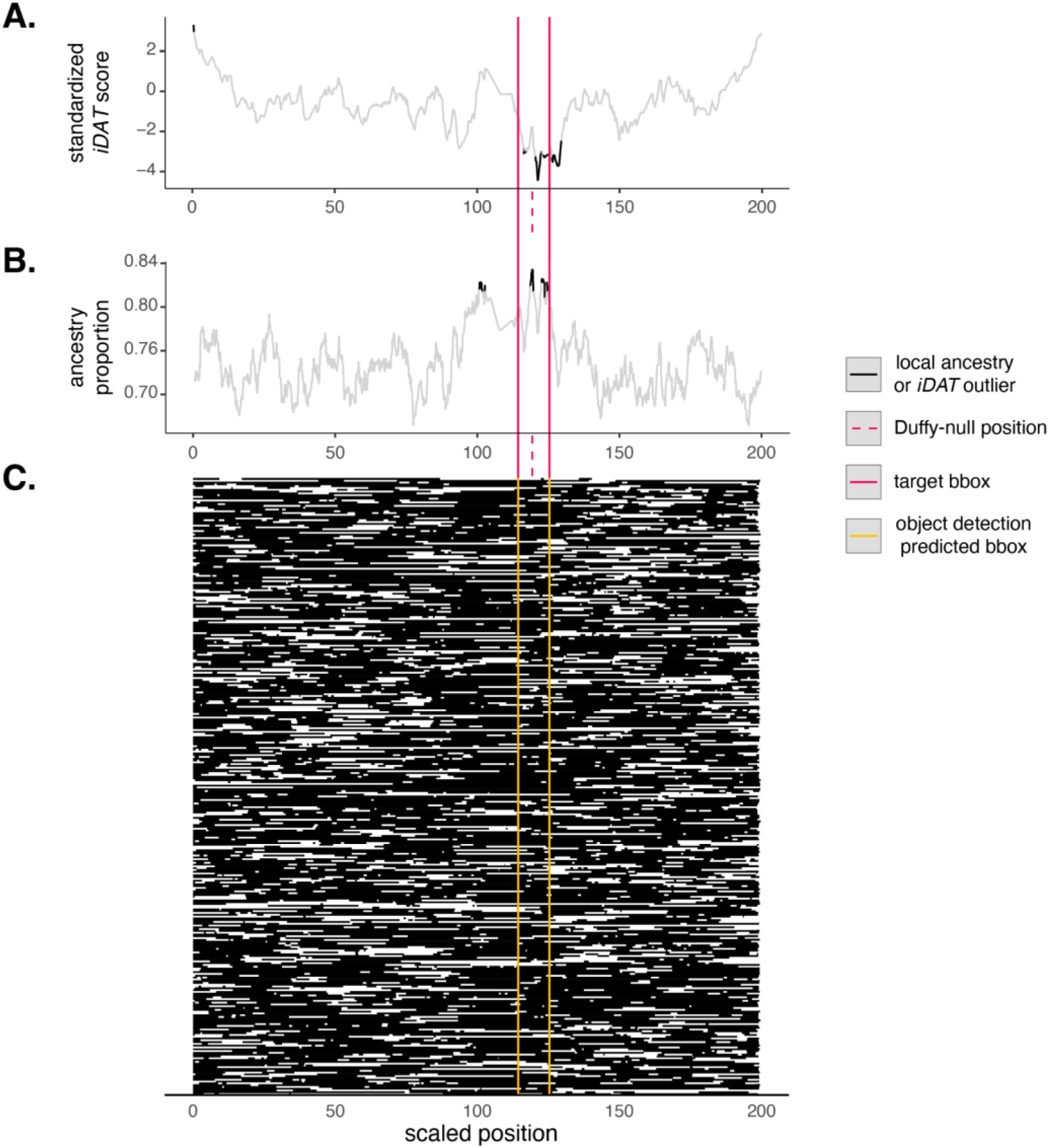
Identification of a known adaptive allele in a human population using multiple ancestry-based methods. We compare multiple methods to detect a well-known example of post-admixture positive selection in the admixed human populations from Santiago, Cabo Verde on the Duffy-null allele protective against *P. vivax* malaria (Hamid et al., 2021). (A) *iDAT* from Hamid et al., 2021, (B) ancestry outlier detection using a 3 standard deviation cutoff, and (C) the object detection approach developed in this paper. African ancestry in black and European ancestry in white. The image represents the entirety of chromosome 1 for 172 individuals. The dashed line indicates the position of the adaptive allele. The inferred bbox using object detection (C) is in yellow, closely matching the true bbox centered on the adaptive allele (red) in size and location. The other two methods infer multiple and/or longer regions as potentially under selection.

Notably, inference was conducted using the pre-trained baseline model whose demographic and genomic scenario differs from that in Cabo Verde. Specifically, the training model included 50% ancestry contributions from each source 50 generations ago; Santiago is estimated to have a 73% African ancestry contribution about 22 generations ago (Hamid et al., 2021; Korunes et al., 2022). We also trained on a 50 Mb window and applied the method to the whole ~250 Mb chromosome 1. Despite these substantial differences, the method performs well, suggesting it can be used widely for populations without well-studied demographic histories. Further, leveraging the general applicability of the baseline model, we made the pre-trained baseline model available online at https://huggingface.co/spaces/imanhamid/ObjectDetection_AdmixtureSelection_Space (see Data and Code Availability). Users can upload an image of painted chromosomes and quickly use the pre-trained set to get inferred adaptation under our method.

In this example, we used genetic recombination distance rather than physical distance. To consider how this choice impacts inference, we generated an image from the Cabo Verde ancestry calls for chromosome 1, but we used physical distance rather than genetic distance. Then, we uploaded that image to the online app with the pretrained data. The model predicts a single bounding box corresponding to physical positions 134,370,749 - 148,191,519. For reference, Duffy-null is at physical position 159,174,683 in this genome build (GRCh37). The center of the bbox is ~18Mb away from Duffy-null. This suggests longer tracts spanning the centromere are affecting the model’s ability to localize the selection signal surrounding the Duffy-null allele. That is, when using physical distance, the model detects a region nearby but less localized to a site under selection, likely owing to recombination interference from the centromere. Therefore, for the purpose of applying this method to real data, users can consider training a model using relevant recombination maps for their system. Alternatively, for reasonably strong performance, users can do as we did here, and generate images using genetic map when inferring with a model that was trained using a uniform recombination map.

## Discussion

We developed a deep learning object detection strategy to detect and localize within the genome post-admixture positive selection based on images of chromosomes painted by local genetic ancestry. Our results demonstrate the power gained when including spatial patterns of ancestry beyond single locus summary statistics, and emphasize the need for further development of methods tailored to populations that do not fit the expectations of classical population genetics methods.

Our object detection approach can leverage complex local ancestry patterns without discarding information about the surrounding genomic context or requiring user choice of statistics. Using simulated and empirical human genetic data from Cabo Verde, we show that our framework better localizes the adaptive locus to a narrower genomic window and is less prone to false positives compared to common ancestry outlier approaches (Figures 3, 4). We expect many empirical examples to actually perform better than this case study because admixture is so recent (~22 generations) and strong (*s* = 0.08 from Hamid et al., 2021) with ~73% admixture contributions from the source with the adaptive allele, which together produce extremely long stretches of African ancestry often spanning the entirety of chromosome 1 reminiscent of the poor performance observed in Table 2 for *m* = 0.9. In both simulated data and our empirical example, the object detection approach remains generally effective at identifying selection even when we misspecify aspects of demographic history such as admixture proportion, admixture timing, and population size trajectory. That is, we expect strong performance on empirical data even without knowing the full details of an admixed population’s history. The size of the window that our method identifies will depend on the chromosome size, input image size, and choice of bbox size used in training. It may indeed be possible to identify a narrower window for a small chromosome, a larger image, or if we train with smaller target boxes. The midpoint of the bbox is a reasonable metric for a point estimate for the location of the adaptive locus.

Despite the overall strong performance of the method, we note several potential pitfalls and areas where future work could make this type of approach more generalizable. A primary barrier to effective implementation is the availability and accuracy of local ancestry calls. As with all ancestry-based approaches, such as ancestry outlier scans, local ancestry calling is a necessary prerequisite for this method. Many tools exist to infer local ancestry along admixed chromosomes, including recent developments for samples in which it is difficult to confidently call genotypes because of low or sparse coverage (Schaefer et al., 2016; Schaefer et al., 2017; Schumer et al., 2020; Wall et al., 2016). Still, local ancestry calling remains potentially challenging, especially in nonmodel systems, and the quality of local ancestry estimates often depends on reference dataset availability and the degree of differentiation between source populations. Notably, we tested our object detection strategy using phased ancestry haplotypes, and further work is needed to address the effects of phase errors. Phasing accuracy can be sensitive to factors such as the availability of reference panels, the number of unrelated individuals present in the sample, and the choice of phasing method (Browning & Browning, 2011). The extent of the impact will vary by species, and empirical tests suggest phasing error is minor in humans (Belsare et al. 2019). The pixel structure that combines multiple loci per pixel may smooth over some of the impact of errors at short stretches of base pairs. We recommend that researchers hoping to take ancestry-based approaches to detecting selection first confirm the validity of their local ancestry calls, for example by first simulating admixed haplotypes from genomes representing proxies for source populations and testing local ancestry assignment accuracy (Schumer et al., 2020; Williams, 2016). Though local ancestry calling is necessary, the similar performance of the object detection method in the high resolution and low-resolution ancestry scenarios demonstrates the utility of our method for a variety of organisms or situations where a limited set of markers are available for assigning local ancestry. Compared to local ancestry outlier approaches, our method may include a potential loss of information or resolution from binning many sites into much fewer possible pixels. However, the selected locus is unlikely to be near the edge of an ancestry tract, and we focus on selection within the last ~100 generations or less; therefore, we expect tracts to be quite long and regions prone to binning error (i.e. edges) constitute a small proportion of the overall tract length. If resolution is a concern, researchers can consider testing different image sizes or genomic window sizes as well.

Ancestry-based methods such as the one presented here that leverage long stretches of higher than expected frequency are well-suited to detect selection on short timescales; we focus on history within a couple hundred generations after admixture and selection onset. For admixture more than a few hundred generations old, the length of ancestry tracts will decay due to recombination over time. As local ancestry at distant sites is decoupled over generations, detectable signatures of long ancestry tracts or high ancestry proportion in a large genomic region surrounding a variant under selection are less likely. Therefore, ancestry-based approaches are better suited for detecting post-admixture selection on the scale of tens to hundreds of generations since admixture. The optimal detection time frame (in generations) will depend both on strength of selection and the timing and proportion of admixture. When admixture is older, assuming selection occurs immediately post-admixture, there has been more time for ancestry tract lengths and frequencies to diverge between neutral and selected sites. That is, recombination has had time to break up ancestry tracts in neutral regions, while the ancestry tracts remain longer in the selected region. So, ancestry-based methods such as ours may perform slightly better for older admixture scenarios (Table 2 & Figure S1). However, this increase in accuracy is true only until a point: if enough time has passed or the selected allele has fixed, the haplotypes decay such that detection of sites under selection becomes more difficult.

Many of the methods we consider in this study, including the object detection method presented here, use the length of ancestry tracts to detect selection. This signature is influenced by the recombination landscape. We demonstrated the impact of one type of recombination nonuniformity, centromere interreference, in the empirical example from Cabo Verde. Notably, the impact was different for the common local ancestry outlier approach, *iDAT*, and our object detection method. Local ancestry outlier approaches may have increased false positives and poorer localization if selection occurs in a low recombination region as local ancestry proportions are impacted at wider distances surrounding a selected variant. The recombination landscape will also affect *iDAT* because the statistic is based on the length of tracts in one genomic region compared to others, so the statistic risks both false positives and false negatives when using physical distances. Incorporating genetic map distances into *iDAT* may decrease some of the impact, but this approach has not been tested and may not improve localization. Under the object detection method, if one uses genetic map distances to generate images as done here, the recombination landscape has less of an influence on performance. We further demonstrated this in our example for detecting selection at Duffy-null in Cabo Verde wherein we compared localization using genetic map distances versus physical distance. We saw worse localization using physical distance owing to the nearby centromere decreasing the recombination rate in the region.

Our empirical example also showed the utility of using our pre-trained model available online, even if the model is misspecified. A central choice that users make is the size of the chromosomal window to include in the 200-pixel image. One can consider whole chromosomes, as we did in our empirical example of Cabo Verde, or partial chromosomes, similar to our example with multiple selected sites. In this study, we tested our model on chromosomes ranging from 50-250Mb. Depending on the population, study system, and the size of the chromosomal region included in the image, the 11-pixel bbox will correspond to a different number of SNPs. The ideal size therefore will depend on the study question and selection history of the population, and there may be a tradeoff between the ability to localize a narrower genomic region and the potential loss of information if signatures of selection unable to be captured in too small of a window.

Our empirical example used human genetic data, though post-admixture selection has been observed across a range of organisms. The baseline model scenario is fairly general and not organism specific. For example, the uniform recombination rate used is reasonable for *Anopheles* mosquitoes and humans (though their overall recombination landscapes differ substantially, the mean rate is similar), and the range of chromosome sizes used in inference (50-250Mb) covers a wide range of organisms. However, the accuracy of local ancestry calls may be impacted by the availability of high-quality reference datasets as proxies for source populations. Available references vary by population and organism, so this could preclude applicability of our method for specific study systems.

Our use of out-of-the box object detection frameworks demonstrates that population genetics researchers can apply deep learning applications without prior experience with machine learning techniques. We required only ~1.5 hours to train the object detection method on 8000 images. To train on 800 images, it only took ~15 minutes with comparably high performance (~90% of selected variants detected vs ~95% with more training examples), making optimization and troubleshooting on small training sets possible in a reasonable timeframe before scaling up to a larger final dataset. That is, one may consider using a smaller training set for optimization of window size and other model decisions prior to training on a larger set. Additionally, with the availability of free GPU access via platforms such as Google Colab, deep learning methodology is accessible to researchers without the means or desire to buy their own GPU or pay for access to a remote server. The same training set can be used for multiple regions of the genome and for multiple populations given the limited impact of model misspecification. More generally, the success of our approach suggests that researchers should consider object detection methods for other problems in detecting selection and population genetics.

## Materials and Methods

### Simulations

Simulated data were generated with the forward simulator SLiM 3, combined with tree-sequence recording to track and assign local ancestry (Haller et al., 2019; Haller & Messer, 2019). For our baseline scenario, we considered a single-pulse admixture event between two source populations (Figure 1). One source population was fixed for a beneficial mutation randomly placed along a 50 Mb chromosome, with selection strength drawn from a uniform distribution ranging from 0 to 0.5. The newly admixed population had a population size *N* of 10000, with 50% ancestral contribution from each source. That is, the range of *Ns* is in [0,5000]. Tree sequence files were output after 50 generations. We used a dominance coefficient of 0.5 (an additive model), recombination rate was set to a probability of a crossover of 1.3 ×10^-8^ between adjacent basepairs per gamete. The SLiM script for our baseline model is available on github (https://github.com/agoldberglab/ObjectDetection_AdmixtureSelection/blob/main/admixture.slim)

### Ancestry Image Generation

For each simulation, we used *tskit* to read the tree sequence files and extract local ancestry information for 200 sampled chromosomes from 100 diploid individuals from the admixed population (Haller et al., 2019; Kelleher et al., 2016, 2018). We then used *R* to generate a black and white 200×200 pixel image of the entire set of sampled chromosomes for each simulation (y-axis representing sampled chromosomes, x-axis representing genomic position), with each position colored by local ancestry for that individual chromosome. In these images, “black” represented ancestry from the source population that was fixed for the beneficial mutation, and “white” represented the other source population. That is, each pixel usually contains many sites depending on the length of the chromosome one uses. We chose 200 pixels for convenience, but other sizes could work. Larger images will take up more computational resources for storage and training.

For our high resolution, or full ancestry images, we used true local ancestry at every position. For our low-resolution ancestry images, we used the same simulations but instead only assigned local ancestry at 100 randomly dispersed markers to generate images. We used the same internally consistent markers across all simulations from the same demographic model. This approach to assigning local ancestry allowed us to test the model performance for scenarios where we have only a few Ancestry Informative Markers (AIMs) for population(s) of interest.

### Object detection model architecture and training

We implemented an object detection model using the IceVision computer vision framework (v0.5.2; https://airctic.com/0.5.2/). Specifically, we trained a FasterRCNN model (Ren et al., 2016) (https://airctic.com/0.5.2/model_faster_rcnn/) with the FastAI deep learning framework (built on PyTorch; https://docs.fast.ai/). We used a resnet18 backbone and pretrained model weights from ImageNet (https://image-net.org/).

For the sets of high- and low-resolution ancestry images described above, we generated 8000 images for training and 2000 images for validation from the same demographic model. In object detection models, the goal is to predict a bounding box around an object of interest. Under the IceVision framework, the bounding box is set as [x-min, y-min, x-max, y-max]. In our case, our goal is to detect the position of the selected variant (if there is one). Thus, for each image in our training and validation sets, we defined the target bounding box as an 11-pixel-wide window centered on the selected variant. For example, if the selected variant is in x-axis position 155, the bounding box was defined as [150, 0, 161, 200].

We trained each model for 30 epochs using the *learn.fine_tune* function, freezing the pretrained layers for one epoch. We used a base learning rate of 3 x 10^-3^ and a weight decay of 1 x 10^-2^.

We largely use an out-of-the-box FasterRCNN architecture with preselected hyperparameters; base learning rate & weight decay were based on testing a few different values and picking the one with the best overall performance. Number of epochs was based on the tradeoff between time to train and gain in validation performance.

The high resolution and low-resolution ancestry models were both trained on an NVIDIA GeForce RTX 2080 Ti GPU. The time to train one model was approximately 1.5 hours.

### Bounding box size and genomic resolution

The method can work on other bounding box sizes, however one would need to train a model on their desired bounding box size. As a proof of concept, we retrained a small set (800 training images from our original training set, 200 validation images from our original validation set) to detect bboxes 5 pixels wide, centered on the variant under selection. We then inferred on the remaining 9000 images from our original training and validation sets. We still see reasonably high performance with this smaller bbox size (~86% of variants detected within a bounding box, Precision = 0.768, recall = 0.756) (Supplemental Table 1). Training on more images should improve this performance.

Alternatively, if researchers wanted higher resolution (i.e. narrower windows), it is likely simpler use a smaller chunk of the chromosome to generate images rather than retrain the entire model to your desired window size.

### Detection threshold

The model essentially is performing a classification task that identifies bboxes, and then returns a probability that that bbox actually contains a selected variant. This probability is defined as the bbox score, which can be interpreted as the model’s level of confidence in that predicted bbox. By default, the model will only return a predicted bbox if the score is above 0.5. This is the detection threshold. Users can alter the detection threshold to return bboxes above any arbitrary score (i.e. make the threshold higher if one wants only higher confidence predictions, lower if one wants to increase recall at the risk of lower precision). We used the default detection threshold of 0.5 for all performance evaluations, except in the case of Precision-Recall Curves (and AUC). For those, we calculated PR over a range of 10000 detection thresholds from 0 to 1. Detection threshold can be set during inference by adding the argument to the predict_dl() function in IceVision, or directly in our demo app via the slider input.

### Validation

We evaluated performance on the validation sets using several metrics. We first calculated precision and recall by defining each x-axis pixel position as an independent test. Each image target had 11 true positives (the size of the bbox, ideally centered on the adaptive allele +/- 5 pixels) and 189 negatives. That is, pixels within the true bbox are all labeled as positive and pixels outside the true bbox are labeled as negative. Because some images may have multiple predicted bboxes, and the sizes of these bboxes can vary, the predicted positives and predicted negatives can be greater than or less than 1 for each pixel. For the purpose of getting a single classification for each pixel, if a pixel was predicted within the x-min and x-max of any bounding box with a score above the threshold, it was classified as a “region under selection” (i.e. a “positive” classification). X-axis positions outside all predicted bounding boxes were classified as a “region not under selection” (i.e. a “negative” classification). In this way, we were able to calculate true and false positives and negatives. We defined P-R in this manner to capture multiple aspects of the method’s performance such as how well it identifies a bbox of the correct size in the correct region.

We also defined several other metrics to assist in evaluating object detection performance across different demographic scenarios. First, we calculated the proportion of predicted bounding boxes that contain the true selected variant, which we defined as the bbox detection rate. We chose this metric because some images have more than one predicted bounding box, and some have none. We wanted to correctly punish the model for returning bboxes that did not contain a selected variant. For example, if the model predicts two bboxes for an image, one which correctly contains the selected variant within the bounds, and a second which does not, the method is not performing as well as we would like. A value close to 1 indicates high sensitivity, or that the method is consistently able to detect a region under selection.

We also calculated the average width of the predicted bounding boxes. If the average width is much wider than the 11-pixels we used in training, this may indicate we have low specificity to detect a region under selection. Finally, we calculated the average number of predicted bounding boxes per image. Since we are only simulating one variant under selection, the model should predict 1 bounding box per image. These metrics combined with the more universal precision and recall statistics allowed us to compare performance of our model across different scenarios and between different methods.

Code to calculate metrics during both training and inference is found in our github example notebook (https://github.com/agoldberglab/ObjectDetection_AdmixtureSelection/blob/6fa95b941608292d219585b1bd8b8dec9c315dce/objectdetection_ancestryimages_example.ipynb).

### Model Misspecifications

We tested the performance of our baseline high resolution ancestry model under several demographic model misspecifications (Results & Table 2). For each misspecification scenario, we generated 1000 high resolution full ancestry images (i.e. incorporating full local ancestry information), ran inference using our trained baseline model, and calculated performance metrics detailed in the previous section.

For these simulations, we followed the baseline scenario described previously while changing one feature of the admixture or population history. We tested inference on images generated from different admixture contributions than what we trained on (10%, 25%, 75% or 90% contribution from the source population providing the beneficial mutation), number of generations since admixture began (25 and 100 generations), population size histories (expansion, contraction, and moderate (50%) and severe (10%) bottlenecks), and a scenario where the selected variant is present in both sources at a frequency of 0.5 (i.e. *F_ST_* of 0 between the sources).

For the population size misspecifications, the expansion (200%) or contraction (50%) events occurred at 25 generations (halfway through the simulation). The bottlenecks occurred at 25 generations and lasted for 10 generations before expanding to the original population size of 10000. All scenarios start with N=10000.

### Comparison to local ancestry outlier approach

We generated 1000 ‘genome-wide’ simulations of 5 independently segregating chromosomes of 50 Mb each. For each simulation, the beneficial allele was fixed at the center of the first chromosome. The rest of the simulation followed exactly the admixture scenario for our baseline model described previously. After sampling 200 haplotypes from the population, we binned the first chromosome into 200 equally-sized windows (to be analogous with the 200×200 pixel images for comparison). Any window with an average local ancestry proportion greater than 3 standard deviations from the genome-wide mean was classified as “under selection” by this outlier approach. We generated ancestry-painted images from the same simulated chromosomes and classified regions under selection using our object detection method trained on the baseline high resolution ancestry scenario.

### Application to human SNP data from Cabo Verde

We used local ancestry calls for ~800k genome-wide SNPs from a previous study of post-admixture selection in Cabo Verde, which included 172 individuals from the island of Santiago (Beleza et al., 2013; Hamid et al., 2021). We focused on Santiago because we had previously detected evidence of strong positive selection in this population for the Duffy-null allele at the *DARC* (also known as *ACKR1*) gene. We generated a 200×200 pixel image of West African and European ancestry tracts on Chromosome 1 for these 172 individuals (344 haplotypes). The length of ancestry tracts can be influenced by the recombination landscape along the chromosome (e.g. long ancestry tracts are often found close to the centromere). To account for this effect, we used genetic map distances rather than physical positions to calculate ancestry tract lengths, and suggest this approach for others using our method if a genetic map is available. We then identified regions under selection on Chromosome 1 using our pre-trained high resolution object detection method for the baseline ancestry scenario (Figure 4).

To compare our results to the local ancestry outlier approach, we identified sites where the proportion of individuals with West African ancestry was more than 3 standard deviations from the mean genome-wide ancestry proportion (~0.73).

We also compared our results to the calculated *iDAT* values from Hamid et al. 2021 (the full genome-wide *iDAT* scores can also be downloaded from Hamid et al.’s associated github repository). This data consists of *iDAT* values for 10,000 randomly sampled SNPs across the genome. *iDAT* is a summary statistic designed to detect ancestry-specific post-admixture selection by calculating the difference in the rate of tract length decay between two ancestries at a site of interest, similar to how *iHS* compares the decay in homozygosity between haplotypes bearing the ancestral and derived alleles at a focal site (Voight et al., 2006). Duffy-null was previously shown to be in a genomic window with extreme values of *iDAT* in Santiago, indicative of the strong recent positive selection at the locus. For our purposes, we first standardized *iDAT* by the genome-wide background. Then, we identified standardized *iDAT* values on Chromosome 1 that were more than 3 standard deviations from the mean genome-wide standardized *iDAT*.

## Acknowledgements

This work was supported by National Institutes of Health R35GM133481 to AG, R35GM138286 to DRS, and F32GM139313 to KLK. We thank Alejandro Ochoa for valuable feedback. We thank Hua Tang and Greg Barsh for generating genetic data used in this study, and the individuals from Cabo Verde for their participation.

## Data and Code Availability

Code for this study is available at https://github.com/agoldberglab/ObjectDetection_AdmixtureSelection. The pretrained high resolution baseline model that was used for most analyses in this study is uploaded and deployed at https://huggingface.co/spaces/imanhamid/ObjectDetection_AdmixtureSelection_Space. Here, users can input a 200×200 pixel, black and white, ancestry-painted image and the model will return vertices and scores for bboxes centered on predicted regions under selection (if there are any). We recommend that users follow the example code in our github for generating ancestry images to ensure that files are in the correct format. We emphasize that this model is trained under a simple single-locus selection scenario, so users should use discretion when deciding if this is an appropriate method for their data. Inferred local ancestry information for the individuals from Cabo Verde can be found at https://doi.org/10.5281/zenodo.4021277, originally published by Hamid et al. 2021 from genotype data published in Beleza et al. 2013.

---

**Supplemental Figure 1.**
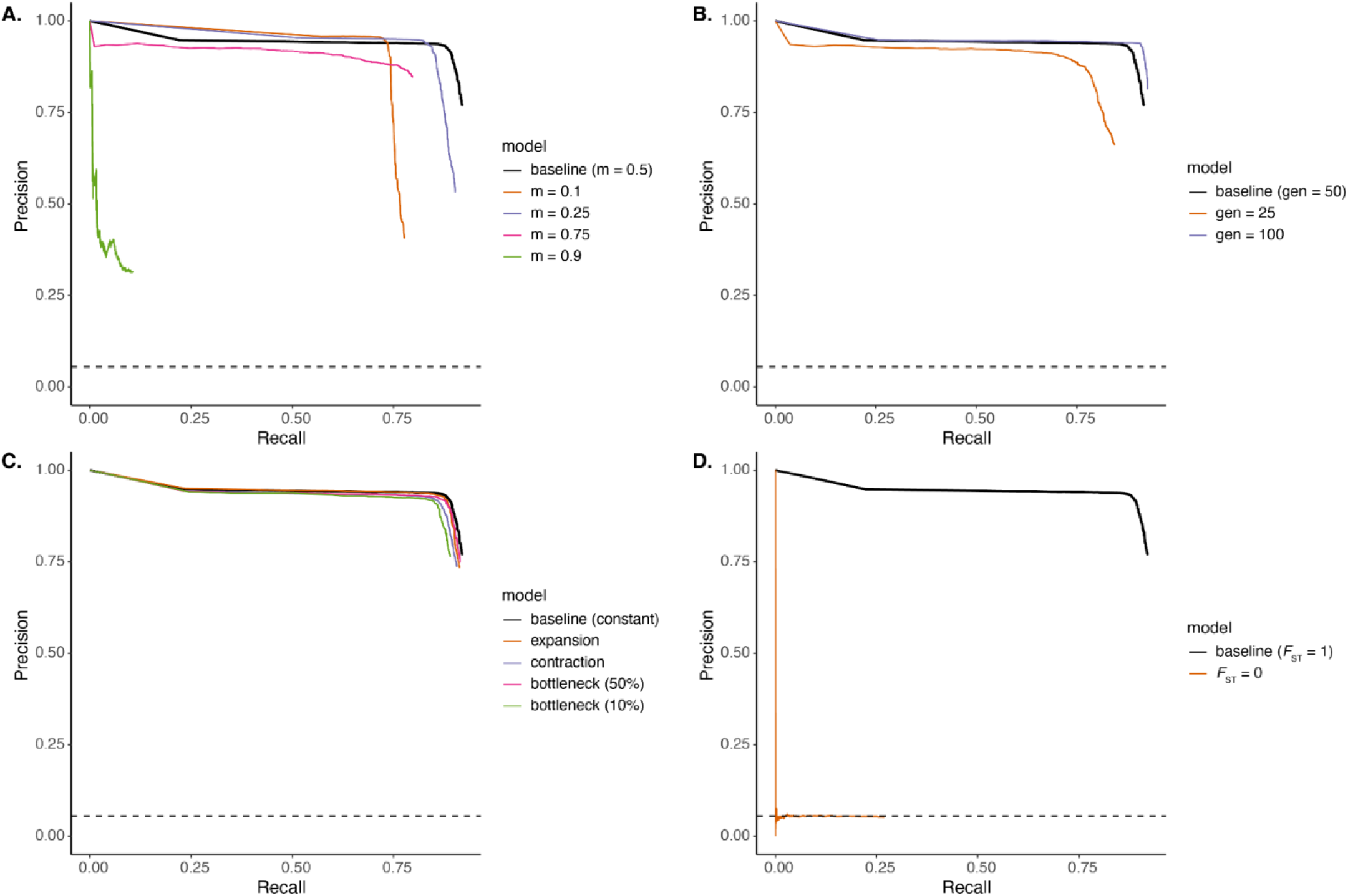
Precision-Recall curves comparing performance under demographic model misspecifications to the baseline scenario for high resolution full ancestry images; baseline is the solid black line in each plot. Panels show different categories of misspecification: A) founding admixture contribution from the population providing the beneficial allele, B) number of generations since admixture occurred, C) population size change since the founding of the admixed populations, and D) level of differentiation between the source populations for the variant under selection. Area under the curves (AUC) can be found in Table 2. The no-skill classifier is indicated by the dashed black lines in each plot.

**Supplemental Figure 2.**
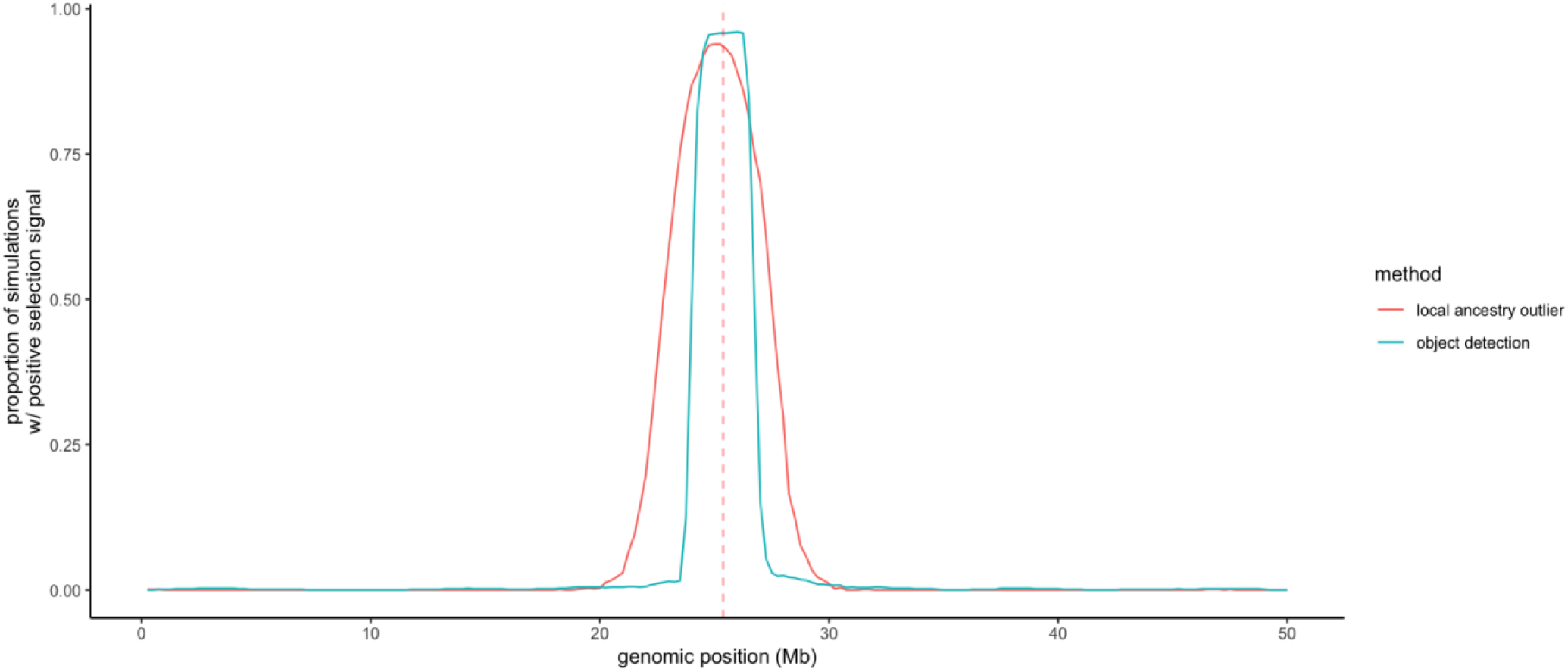
Comparison of local ancestry outlier approach and object detection method. Replot of data from Figure 3A, showing, for each genomic window, the proportion of simulations that had that region classified as “under selection” by either the object detection or local ancestry outlier methods.

**Supplemental Figure 3.**
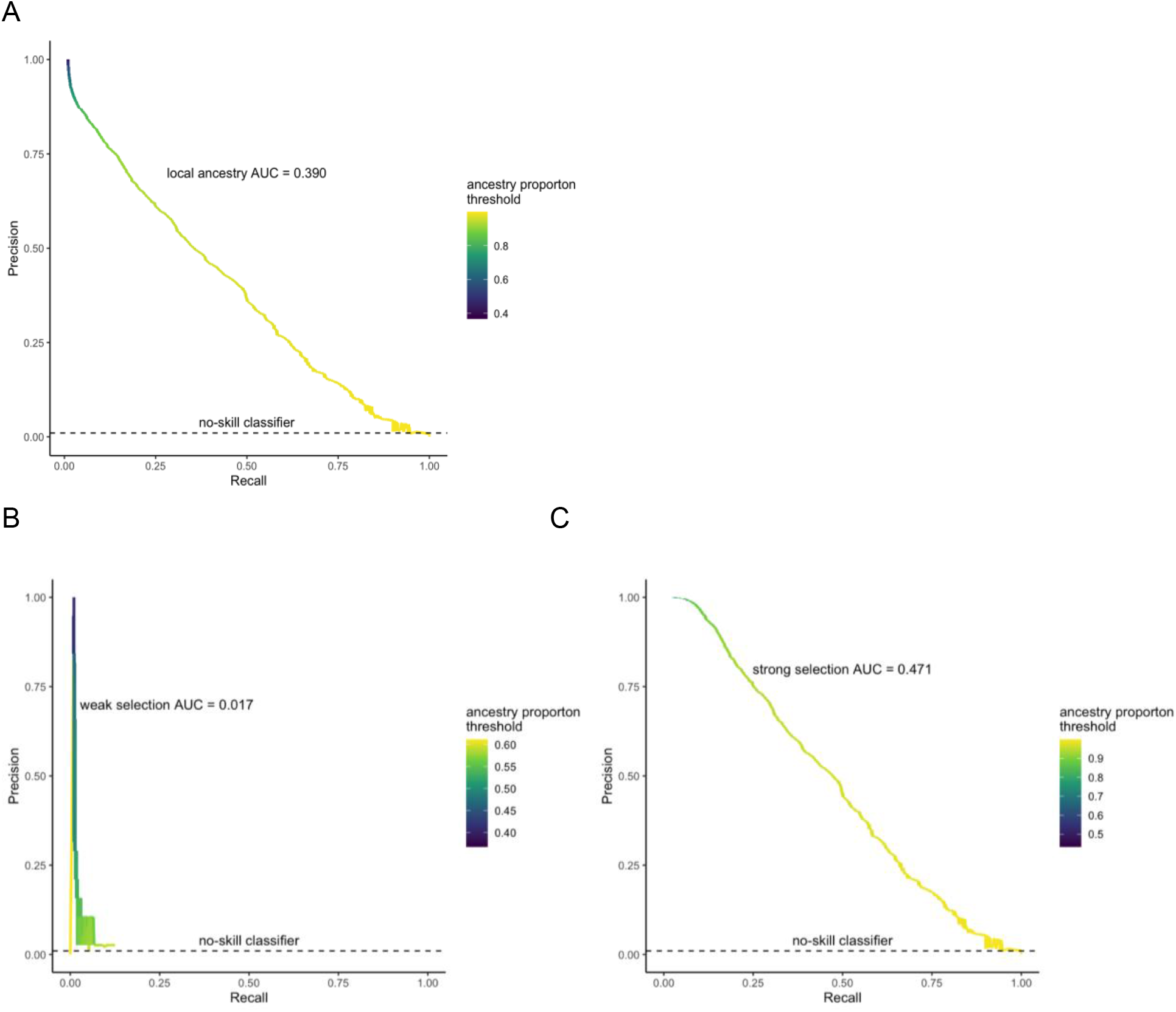
Alternative measure of performance of local ancestry outlier approach. We used the same simulations that were generated for Figure 3 over a range of selection coefficients. We defined the “prediction score” as the ancestry proportion, and calculated PR over the range of local ancestry proportions (~0.367 to ~1). Because the “selected variant” is at the very edge of the 100th window, we labeled both windows 100 and 101 as “positives” and everything else as negatives. (A) across selection coefficients. (B) Splitting into “weak selection” simulations (s < 0.01, n = 3800 [200 windows for 19 simulations]) and (C) “strong selection” simulations (s > 0.1), n = 162600 [200 windows for 813 simulations]). Evaluating performance in this way punishes the local ancestry method more than Figure 3 because the wide affected region with high ancestry proportion results in low recall over a range of outlier “thresholds.”

**Supplemental Figure 4.**
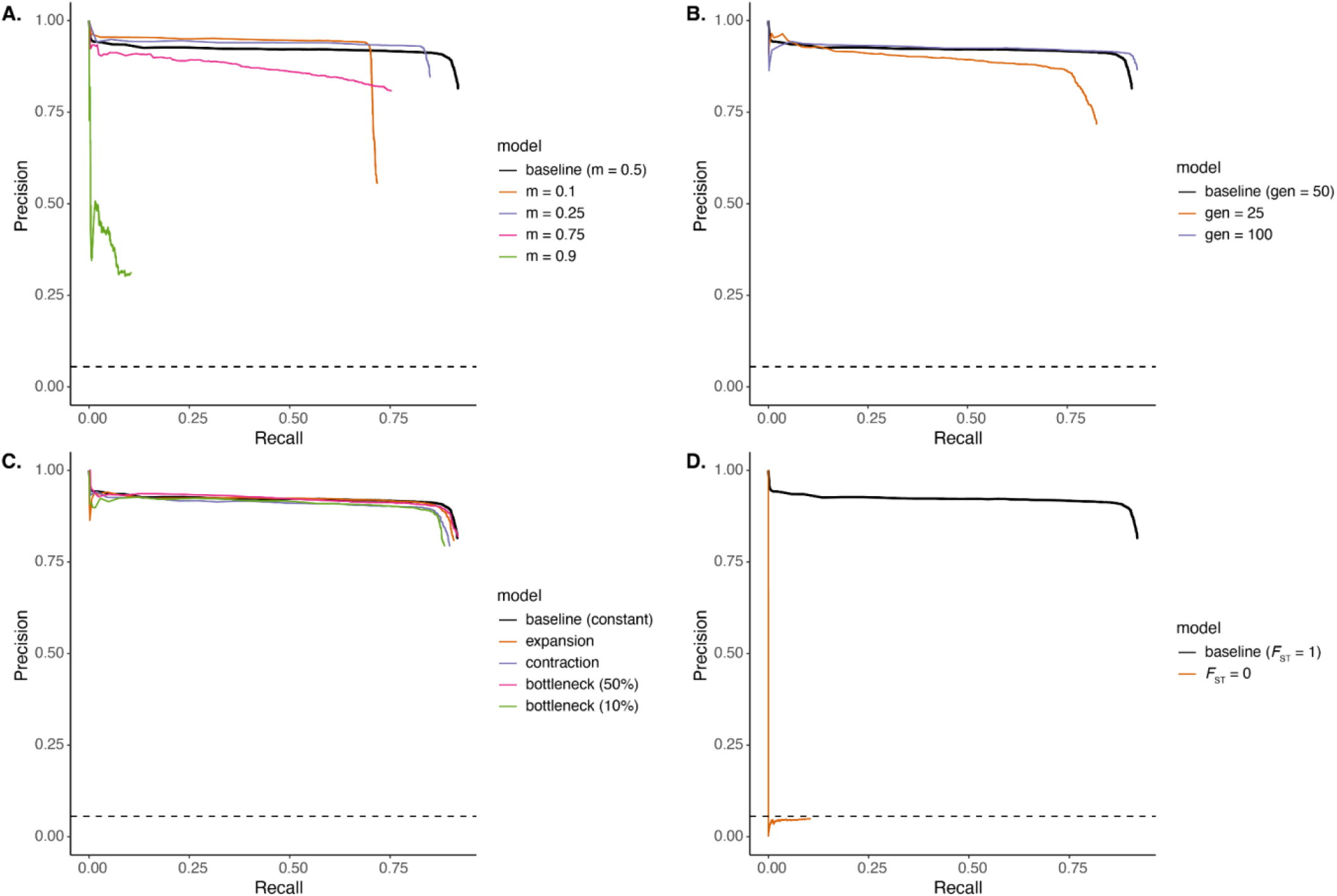
Precision-Recall curves comparing performance under demographic model misspecifications to the baseline scenario (i.e. the scenario that the network was trained on) for low-resolution ancestry resolution images; baseline is the solid black line in each plot. Panels show different categories of misspecification: A) founding admixture contribution from the population providing the beneficial allele, B) number of generations since admixture occurred, C) population size change since the founding of the admixed populations, and D) level of differentiation between the source populations for the variant under selection. Area under the curves (AUC) can be found in Table S2. The no-skill classifier is indicated by the dashed black lines in each plot. Analogous to Figure S1 for high-resolution ancestry.

**Supplemental Table 1.** Performance of object detection method with a smaller 5-pixel bbox using 800 training images and 200 validation images.

| ancestry resolution | bbox detection rate | average width | average number of bounding boxes | precision | recall |
| --- | --- | --- | --- | --- | --- |
| high (full ancestry) | 0.861 | 4.956<br>(var: 0.264,<br>n=8561) | 1.033<br>(var: 0.160,<br>n = 9000) | 0.768 | 0.756 |

**Supplemental Table 2.**
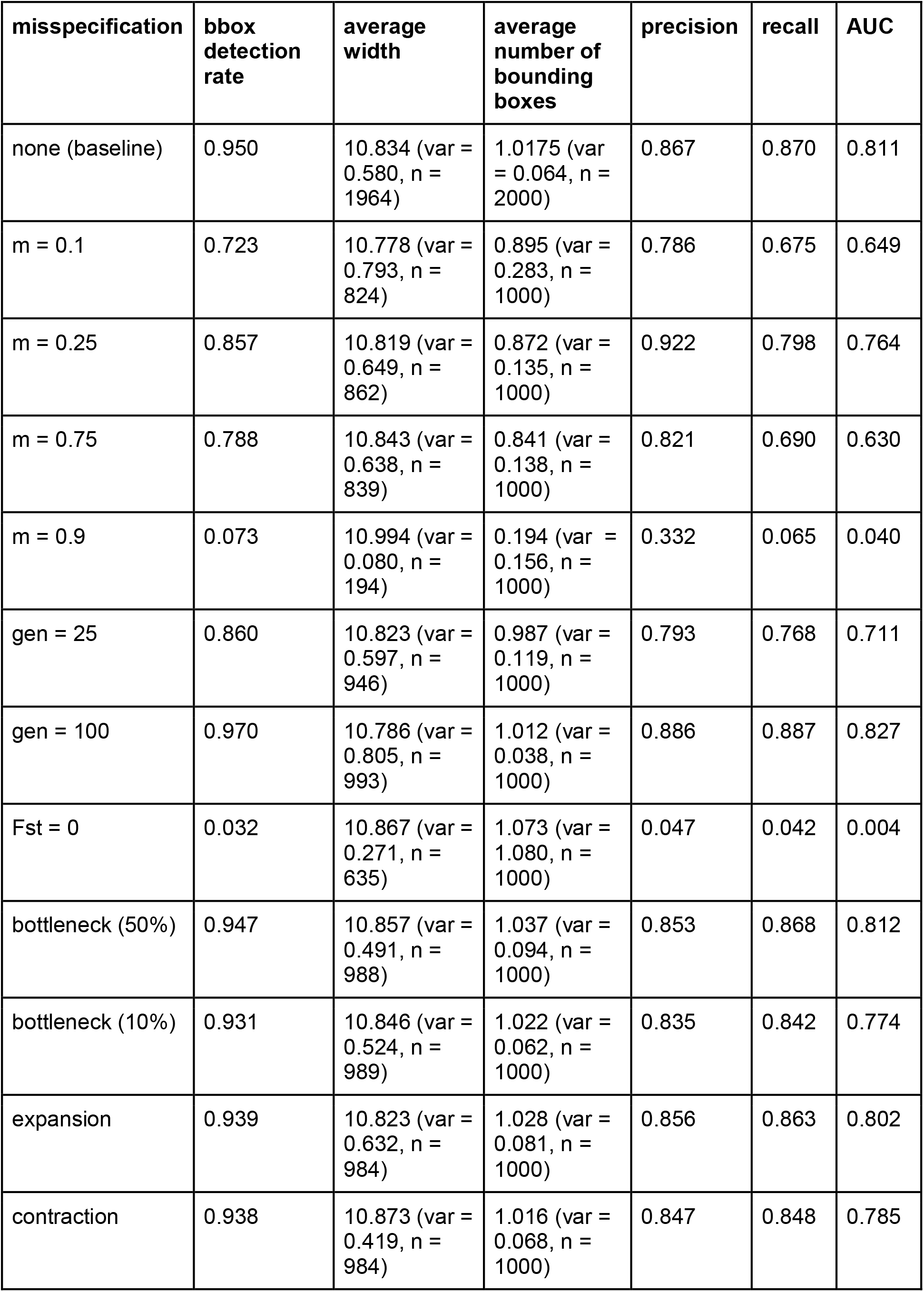
Performance of object detection method on images generated from demographic misspecifications for low resolution ancestry. Further details of models in Materials and Methods, Figure S4.

| misspecification | bbbox<br>detection<br>rate | average<br>width | average<br>number of<br>bounding<br>boxes | precision | recall | AUC |
| --- | --- | --- | --- | --- | --- | --- |
| none (baseline) | 0.950 | 10.834 (var = 0.580, n = 1964) | 1.0175 (var = 0.064, n = 2000) | 0.867 | 0.870 | 0.811 |
| m = 0.1 | 0.723 | 10.778 (var = 0.793, n = 824) | 0.895 (var = 0.283, n = 1000) | 0.786 | 0.675 | 0.649 |
| m = 0.25 | 0.857 | 10.819 (var = 0.649, n = 862) | 0.872 (var = 0.135, n = 1000) | 0.922 | 0.798 | 0.764 |
| m = 0.75 | 0.788 | 10.843 (var = 0.638, n = 839) | 0.841 (var = 0.138, n = 1000) | 0.821 | 0.690 | 0.630 |
| m = 0.9 | 0.073 | 10.994 (var = 0.080, n = 194) | 0.194 (var = 0.156, n = 1000) | 0.332 | 0.065 | 0.040 |
| gen = 25 | 0.860 | 10.823 (var = 0.597, n = 946) | 0.987 (var = 0.119, n = 1000) | 0.793 | 0.768 | 0.711 |
| gen = 100 | 0.970 | 10.786 (var = 0.805, n = 993) | 1.012 (var = 0.038, n = 1000) | 0.886 | 0.887 | 0.827 |
| Fst = 0 | 0.032 | 10.867 (var = 0.271, n = 635) | 1.073 (var = 1.080, n = 1000) | 0.047 | 0.042 | 0.004 |
| bottleneck (50%) | 0.947 | 10.857 (var = 0.491, n = 988) | 1.037 (var = 0.094, n = 1000) | 0.853 | 0.868 | 0.812 |
| bottleneck (10%) | 0.931 | 10.846 (var = 0.524, n = 989) | 1.022 (var = 0.062, n = 1000) | 0.835 | 0.842 | 0.774 |
| expansion | 0.939 | 10.823 (var = 0.632, n = 984) | 1.028 (var = 0.081, n = 1000) | 0.856 | 0.863 | 0.802 |
| contraction | 0.938 | 10.873 (var = 0.419, n = 984) | 1.016 (var = 0.068, n = 1000) | 0.847 | 0.848 | 0.785 |

